# Elucidating the roles of Alzheimer disease-associated proteases and the signal-peptide peptidase-like 3 (SPPL3) in the shedding of glycosyltransferases

**DOI:** 10.1101/317214

**Authors:** Assou El-Battari, Sylvie Mathieu, Romain Sigaud, Maëlle Prorok-Hamon, L’Houcine Ouafik, Charlotte Jeanneau

## Abstract

The Golgi resident glycosyltransferases (GTs) are membrane-bound glycoproteins but are frequently found as soluble proteins in biological fluids where their function remains largely unknown. Previous studies have established that the release of these proteins involved Alzheimer disease-associated proteases such as β-secretases (BACE1 and BACE2) and the intramembrane-cleaving aspartyl proteases Presenilins 1 and 2. Recent studies have involved another intramembrane-cleaving enzyme, the signal peptide peptidese-like-3 (SPPL3). Except for the latter, the two former studies mostly addressed particular cases of GTs, namely ST6Gal-I (BACEs) or GnT-V (Presenilins). Therefore the question still remains as which of these secretases is truly responsible for the cleavage and secretion of GTs. We herein combined the 3 proteases in a single study with respect to their abilities to release 3 families of GTs encompassing three N-acetylglucosaminyltransferases, two fucosyltransferases and two sialyltransferases. Green fluorescent protein (gfp)-fused versions of these GTs were virally transduced in mouse embryonic fibroblasts devoid of BACEs, Presenilins or SPPL3. We found that neither BACE nor Presenilins are involved in the shedding of these glycosyltransferases, while SPPL3 was involved in the cleavage and release of some but not all GTs. Notably, the γ- secretase inhibitor DFK-167 was the only molecule capable of significantly decreasing glycosyltransferase secretion, suggesting the involvement of γ-secretase(s), yet different from Presenilins but comprising SPPL3 among other proteases still to be identified. Using confocal microscopy, we show that SPPL3 selectivity towards GTs relays not only on sequence specificity but also depends on how GTs distribute in the cell with respect SPPL3 during their cycling within and outside the Golgi.

## INTRODUCTION

Cell surface glycans participate in a multitude of cellular processes including the immune cell trafficking [1], autoimmunity and cancer [2], neural development and plasticity [3] as well as interactions between bacteria and epithelial cells in the gastrointestinal system [4], to name a few. To build-up these structures the glycosyltransferases act between sugar donors and acceptors in a concerted and regulated manner. The biosynthesis of *N*- glycoproteins for example, requires multiple steps and multiple cellular compartments starting in the endoplasmic reticulum (ER) where glycosylation is initiated, then transiting the Golgi apparatus where glycans are edited/elongated/modified, before moving to their final destinations [reviewed in [5]]. Golgi glycosyltransferases (GTs) have a type-II orientation as they contain a short cytosolic amino-terminal tail, a single transmembrane domain followed by the so-called stem (or stalk) region and a large carboxy-terminal catalytic domain oriented to the lumen of the Golgi apparatus [reviewed in [6]]. It is generally admitted that glycan structures reflect the sequential order in which glycosyltransfrases are distributed between the *cis*, *medial* and *trans* Golgi cisternae. In addition of being the glycosylation factory of the cell, the Golgi apparatus is also a centerpiece of protein-sorting and secretion. Therefore, glycosylation does not only rely on the expression level of particular GTs or on the way they segregate into distinct Golgi compartments, but also on how their steady-state is maintained between retention and secretion mechanisms [7]. Furthermore, although targeting and retention of GTs into specific Golgi compartments mainly depend on the cytosolic and transmembrane domains [8], their steady-state localization seems to be influenced by dynamic processes involving iterative cycles within the Golgi and between the Golgi and the endoplasmic reticulum [9,10].

On the other hand, some glycosyltransferases are able to oligomerize through their transmembrane and/or stem domains to form large aggregates that favor their retention in respective cisternae [10], while some others are shed from cells by proteolytic cleavage and found as truncated proteins in culture media and body fluids [11]. In this regard, studies on secreted forms of GalNAcT-I, ST6Gal-I, β4GalT-I and GM2 synthase have revealed that they could be processed at different positions, mostly within the stem region [12,13,14,15,16] but also within the transmembrane domain, as in the case of GnT-V [17]. Other glycosyltransferases such as FucT-I or FucT-VII were not found in cell culture media [18,19]

The release of soluble glycosyltransferases from cells and its biological significance still raises several issues. Saito and colleagues have shown that the β1,6-N-acetyl-glucosaminyltransferase-V (GnT-V), released from human colon carcinoma WiDr cells, is involved in tumor angiogenesis in a glycosylation-independent manner [20]. Later on, Nakahara *et al.* [17] reported that the shedding of this angiogenesis inducer is initiated by the intramembrane protease complex Presenilin, thus suggesting that there might be more than one protease involved in the release of glycosyltransferases. So far, most of the studies cited above have been carried out on one single glycosyltransferase at a time, using approaches and cell systems often different from one laboratory to another; which in turn hampers the generalisation of the data. More recently Fluhrer and collegues elegantly demonstated that the shedding of glycosyltransferases is rather mediated by the signal peptide peptidase SPPL3, the loss of which directly influenced the glycosylation state of glycoprotein substrates [21]. In a further study, the authors used a proteomic approach, the so-called «secretome protein enrichment with click sugars (SPECS)» to screen for other candidates cleaved by SPPL3, including GTs [21]. These studies clearly showed that all SPPL3 substrates are type-2 glycoproteins [21]. Yet, SPECS data showed some discrepancies with experimental models including cells lacking SPPL3 or those overexpressing this protease. This is notably the case of ST6Gal-I and to some point, that of GnT-V; both GTs being reported by others as substrates of the Alzheimer’s disease-associated proteases BACE-1 and Presenilin-1, respectively [17,22].

Therefore, to address the issue of which protease is actually involved in glycosyltransferase secretion, it becomes highly desirable to carry out a comparative study using a unique cell line expressing a set of GTs, in which a given protease gene is present or suppressed. We therefore herein used the mouse embryonic fibroblasts (MEF cells) lacking genes of either BACEs or Presenilins, as well as those in which the SPPL3 gene was deactivated by a CRISPR/Cas9 approach. We analyzed the release of several glycosyltransferases, including ST6Gal-I and GnT-V, and found that they are all are secreted from cells and recovered in culture media, regardless of BACE(s) or Presenilin(s). SPPL3 however, is required for the release of some, but not all glycosyltransferases.

## MATERIALS AND METHODS

### Cells

Wild-type (WT) mouse embryonic fibroblasts (MEFs) and those originating from BACE1 KO mice (referred to as B1-KO cells) or BACE1 and BACE2 double KO mice (referred to as B1/2-KO cells) were generated as described previously [23]. MEFs from Presenilin-1 and Presenilin-2 double KO mice (referred to as PS-dKO) were produced as described [24,25]. All those cell lines were kindly provided by Dr. De Strooper (Katholieke Universiteit, Leuven, and Flanders Institute for Biotechnology, Belgium). The SPPL3-knockout MEFs cells were generated by CRISPR/Cas9 genome editing according to Mali [26] (see below for details). MEFs cells were cultured in Dulbecco’s modified Eagle’s medium (DMEM) containing 10% fetal calf serum (FCS), 100 U/ml penicillin and 100 μg/ml streptomycin. The human embryonic kidney cells HEK-293T were maintained in the same medium as above. All culture reagents were from Invitrogen (Cergy-Pontoise, France).

### DNAs, CRISPR/Cas9 constructs and lentiviral transduction

Except for GnT-V, all other GTs were used as C-terminally gfp-tagged proteins as previously described [19]. In the case of GnT-V, a FLAG-tagged version was preferred to a gfp-tagged one because of the unusual large size (~100 kDa) of this glycosyltransferase [27] compared to other GTs (~50 kDa). To this end, the FLAG sequence (GATTACAAGGATGACGACGATAAG) was fused to the 3’ end of GnT-V cDNA, avoiding the stop codon, by overlapping PCR and the resulting DNA was subcloned between *Age*I and *Sal*I sites of the the lentiviral vector pRRLSIN.cPPT.PGK-GFP.WPRE (referred to as pL1, kindly provided by Dr. Trono, University of Geneva, Switzerland), giving rise to pL1/GnT-V-FLAG construct. The chimeric β1,3GnTIII/FucTVII-gfp was made by fusing the first 29 amino acid sequence of β1,3GnT-III (encompassing the cytosolic and the transmembrane domain) to the catalytic domain of FucT-VII (starting at Ser35), essentially as described [8]. The mCherry-tagged murine SPPL3 (mCherry-SPPL3/pcDNA3) [28], was kindly provided by Dr. Pomerantz (The Johns Hopkins University School of Medicine, Baltimore, Maryland, USA). To generate SPPL3-knockout MEF cells by CRISPR/Cas9 genome-editing technology, an SPPL3-specific gRNA-encoding the sequence 5’-ATCGGGGACATTGTGATGCC-3’ was chosen according to [29] and verified for absence of off-targets using the Optimized Crispr Design software (https//:crispr.mit.edu). This sequence was cloned in the *Afl*II site of the gRNA cloning vector (Addgene #41824) according to Mali *et al*., [26]. The sgRNA cassette was then PCR ampified and subcloned between *Cla*I and *EcoR*I sites of pLVTHM vector [30], giving rise to pLVTHM/gRNA expression vector. The resulting plasmid was then equiped with Cas9-T2A-mCherry DNA (encompassing Cas9 fused to mCherry through the T2A peptide) to generate an all-in-one pLVTHM/Cas9-T2A-mCherry/gRNA lentiviral system allowing for co-expression of Cas9 and the SPPL3-sepecific gRNA, while easing efficient detection of transduced cells through mCherry staining. For this purpose, Cas9-T2A-mCherry was amplified from the plasmid pU6-(BbsI)_CBh-Cas9-T2A-mCherry plasmid (Addgene #64324) using the Phusion High Fidelity PCR kit (NEB) polymerase and subcloned between the *Pme*I and *Spe*I sites of pLVTHM/gRNA vector. Lentiviral particles were produced by transfecting HEK-293T cells with pLVTHM/Cas9-T2A-mCherry/gRNA construct and packaging vectors as described [31]. Viral-containing supernatants were collected 24, 48 and 72 hours posttransfection, combined and concentrated using the Lenti Concentrator (Origen) according to the manufacturer instructions. Transduction of cells with viral constructs was performed as previously described [31].

### Protein secretion and Western blots

To detect secreted GTs-gfp, culture media were replaced with OptiMEM which was collected after 18-24h incubation at 37°C, cleared from detached cells and debris by centrifugation, filtered through a 0.22-μm pore-size filters and concentrated 10 times by ultrafiltration using Microcon YM-10 concentrator (Millipore, Molsheim France) with 10 kDa cutoff filters. Protein concentrations were determined using the BCA assay (Pierce, Thermo Fisher Scientific, Brebière. France) and samples were normalised accordingly prior to loading on gels. Proteins were then resolved on a 10% SDS/PAGE according to Laemmli [32], transferred to nitrocellulose membranes that were subsequently stained with Ponceau S (0.1% Ponceau S w/v, 5% acetic acid v/v), to confirm equivalent loading and then blocked in 5% milk in TBST overnight. Glycosyltransferases were probed with the anti-gfp mAb JL-8 as described [33], or anti-FLAG M2 mAb (Sigma) to detect the FLAG-tagged GnT-V. SPPL3 was detected with the anti-SPPL3 pAb (Origen, Rockville, USA). Membranes were then incubated with peroxydase-conjugated secondary antibodies and visualized by chemiluminescence using the ECL kit (Amersham Pharmacia Biotech, Buckinghamshire, United Kingdom). Immunoblots shown are representative of at least three independent experiments.

### Inhibitor treatments

Cells expressing GTs were treated with the following inhibitors, the β-secretase inhibitor C3 (BACE inhibitor IV, Calbiochem, Darmstadt Germany) [34], the γ-secretase inhibitor DFK-167 (Enzyme System Products, Livermore CA, USA) [17], the metalloprotease inhibitor GM6001 (Calbiochem). Inhibitor stocks were made in DMSO and added to cells in serum-free OptiMEM medium at 1% final concentration of DMSO and cultured for 18-24h. Culture media that were not treated with inhibitors were supplemented with 1% DMSO. Media were then collected, cleared from cell debris, concentrated 10 fold as described above and subjected to 10% SDS/PAGE.

### Confocal microscopy

For confocal microscopy of living cells, these were seeded on 8 wells chamber slides with poly-Lysine coated coverslips (Lab-Tek, Hatfield PA, USA), 48h before examination. When permeabilization of cells is needed for Golgi imaging, cells were seeded on the same coverslips as above 48h prior to confocal microscopy, fixed with 4% paraformadehyde in calcium/magnesium-free PBS [PBS(-/-)] and permeabilized with 0.5% Triton X100 in PBS(-/-) containing 1% fetal calf serum. Cells were then incubated with goat anti-α-mannosidase-II polyclonal antibody (Abcam, 1/500 dilution) for 1h followed by 15 min in the presence of AlexaFluor-594 labeled rabbit anti-goat antibody (ThermoFisher, 1/500 dilution). Samples were then visualized with a Zeiss CSM410 confocal laser scanning microscope.

## RESULTS AND DISCUSSION

In a previous work, we have expressed six different gfp-tagged glycosyltransferases (GTs-gfp) including β3GnT-III, C2GnT-I, FucT-I, FucT-VII, ST3Gal-I and ST6Gal-I, in CHO cells and found that except for FucT-VII, all GTs-gfp were released in culture media [19]. Others have reported that the cleavage and secretion of ST6Gal-I was due to the Alzheimer’s beta-secretase BACE1 [16,22,35]. Likewise, the gamma-secretase Presenelin-1, another Alzheimer disease-associated protease, has been described as being required for the secretion of the glycosyltransferase GnT-V [17]. Therefore, the present study has become necessary particularly since a recent work has reported that another gamma-secretase SPPL3, is the one responsible for the release numerous glycosyltransferases [36] together with many other type-II glycoproteins.

### Glycosyltransferases are released from BACE1- and BACE2-deficient fibroblasts

In a first set of experiments we sought to verify whether or not BACE1 and BACE2 are involved in the cleavage of GTs, using cells where genes for those proteases have been deleted. For this purpose, we used mouse embryonic fibroblasts derived from BACE1 knock out mice (B1-KO) and BACE1/BACE2 double knock out mice (B1/2-KO) kindly donated by De Strooper Laboratory [23]. These cells were virally transduced with PL1/GTs-gfp and the released proteins were detected on western blots by the anti-gfp mAb JL8. Fig. 1 shows comparison between GTs collected in the media (M) and their intracellular counterparts (C). This figure shows that either FucT-I, ST3Gal-I, ST6Gal-I, C2GnT-I or β3GnT-III, they are all released in culture media whether BACE1 is present (WT) or absent (B1-KO) or when both BACE1 and BACE2 are absent (B1/2-KO). Consistent with our previous finding, FucT-VII was not secreted whatever the cell line considered [19]. These data clearly indicate that deletion of BACE1 and BACE2 genes do not cause any defect in the secretion of the GTs studied herein.

**Figure 1:**
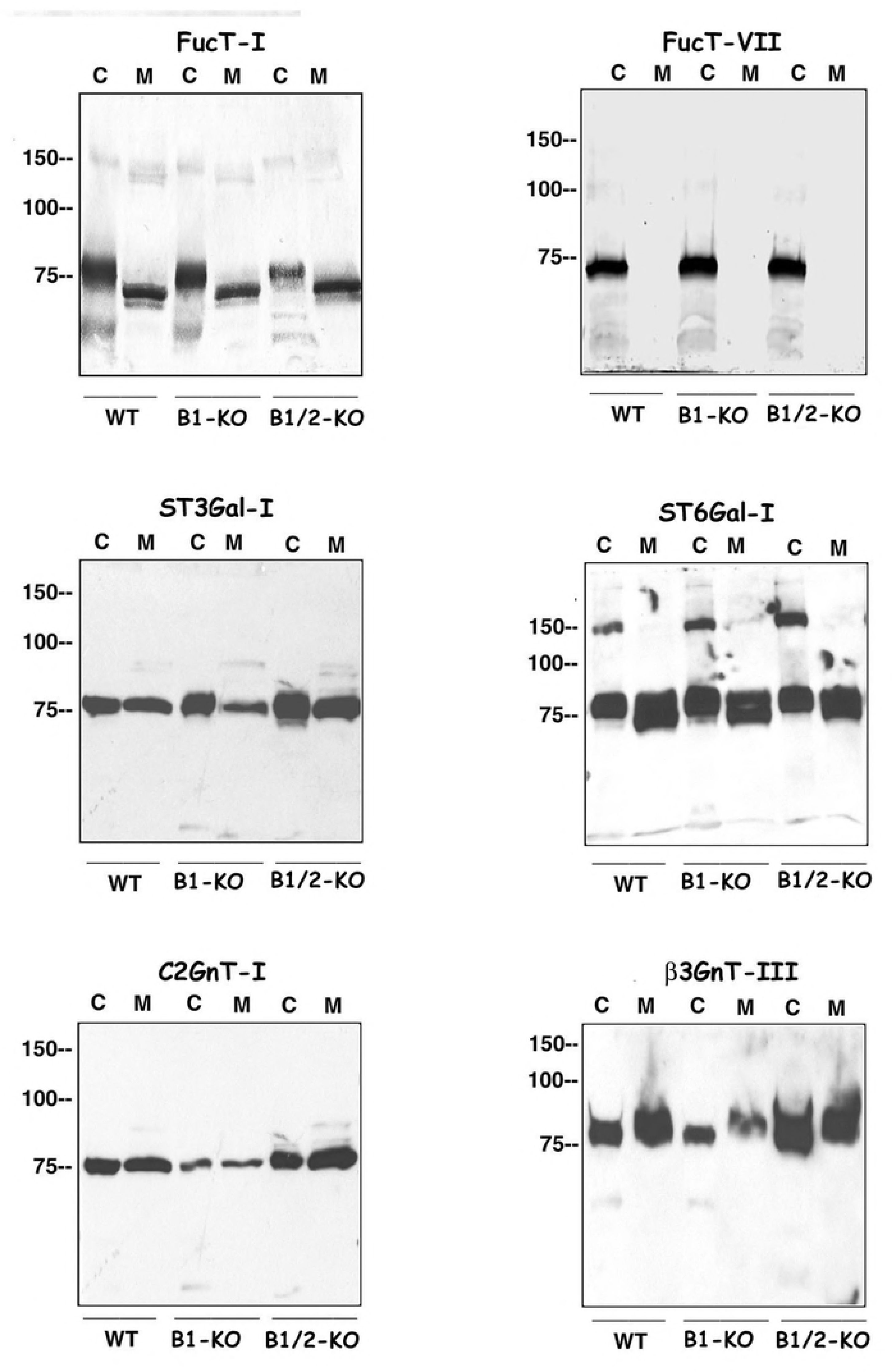
Comparison of GTs secretion in the presence or absence of genes for BACE-1 and BACE-2. Control MEFs (WT) and those where BACE-1 alone or both BACE-1 and BACE-2 genes were suppressed (B1-KO and B1/2-KO, respectively), were transduced with lentiviral pL1 constructs harboring FucT-I, ST3Gal-I, C2GnT-I, FucT-VII, ST6Gal-I or β3GnT-III genes fused to gfp. Culture media were recovered, concentrated as indicated under Material and Methods and protein contents of the samples were determined and normalized in order to load equivalent quantities of material. Proteins were then analyzed by SDS–PAGE and immunoblotting using the anti-gfp mAb JL8. With the exception of FucT-VII which is not found in cell supernatants, all other GTs are released in culture media whether BACE-1 or BACE-2 genes are present (WT) or absent (B1-KO and B1/2-KO).

Considering the sizes of the proteins released in culture media, it appears that many secreted portions of GTs exhibit comparable sizes with their intracellular counterparts (for example C2GnT-I or ST3GalI, see Fig. 1) or even higher molecular weight, such as β3GnT- III (Fig. 1). This could be interpreted as being due i) to a hyperglycosylation of the secreted forms that may compensate the protease-induced lowering of the molecular weight or ii) to a loss of a small protein portion or iii) to a combination of the two situations. We have previously reported that indeed, the secreted β3GnT-III carries more *N*-glycans than its cell-associated counterpart (see Ref 19 and Fig. 7 therein). Now, if hydrolysis occurs within or close to the transmembrane domain of a GT, one would expect that only a small protein stub would be cut-out, leading to a negligible loss in the size of the released protein. To verify this hypothesis we sought to test whether glycosyltransferases could be substrates of the so-called intramembrane cleaving proteases (i-CLiPs). The i-CLiP family includes metalloproteases (S2P), rhomboid serine proteases, signal peptide peptidase aspartyl proteases, and γ-secretase aspartyl protease complexes [37]. In this regard an aspartyl gamma-secretase Presenelin-1, has been implicated in the shedding of the glycosyltransferase GnT-V [17] that belongs to the same family of «branching» glycosyltransferases as C2GnT-I; the former being specific of *N*-glycans [38] while the latter acts exclusively on O-glycans [39]. This therefore prompted us to examine the processing of our GTs under the conditions used by the authors [17], including the use of the γ-secretase inhibitor DFK-167 and the Presenilin1/2-deficient cells.

### The release of glycosyltransferases is impaired by the γ-secretase inhibitor DFK-167

In a first series of experiments, MEF cells expressing ST6Gal-I-gfp were treated with DFK-167 and compared to other inhibitors including GM6001, a broad spectrum α-secretase inhibitor and with the β-secretase inhibitor-IV (C3). As shown on Fig. 2A, the accumulation of ST6Gal-I-gfp in media (M) is not impaired whether the β-secretase (C3) or the α-secretase inhibitors (GM6001) are present or not, while it dramatically decreases in the medium of cells treated with the γ-secretase inhibitor DFK-167 (see also Fig. 2B depicting another experiment where the blot was intentionally overexposed to emphasize the effect of the inhibitor). This result is consistent with previous findings on GnT-V [17] and is indicative of γ-secretases participation in the release of ST6Gal-I-gfp. We then asked how other GTs behave towards this inhibitor. To this end, MEF cells were transduced with different GTs-gfp constructs and treated or not with DFK-167. As shown in Fig. 2C, all GTs exhibit a strongly impaired secretion in the presence of this drug; some even were not secreted at all, such as β3GnT-III or C2GnT-I. This data clearly shows that what is true for GnT-V and ST6Gal-I is true for other GTs and suggest the involvement of DFK-167-sensitive γ-secretase(s) in the this process.

**Figure 2:**
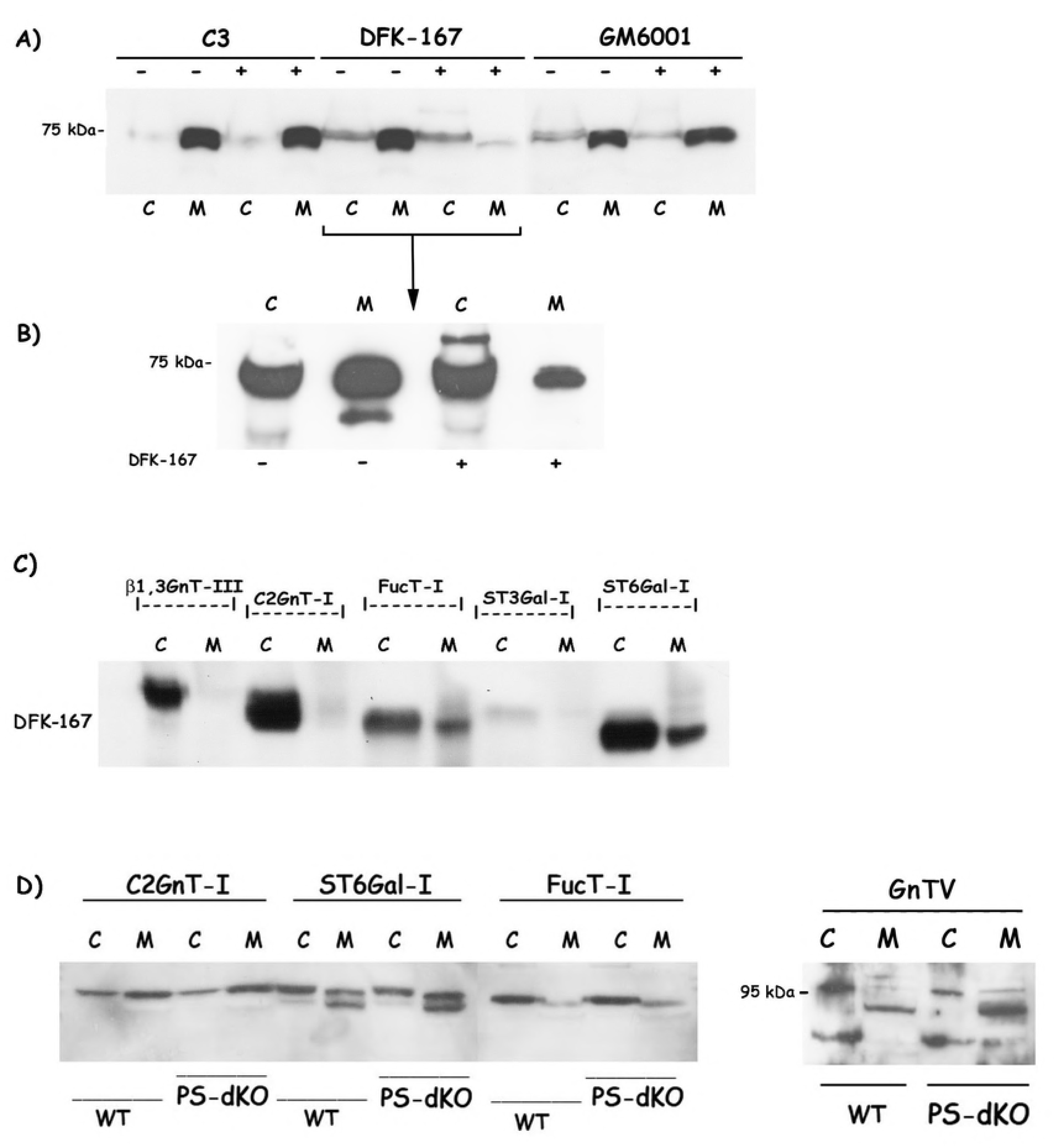
Glycosyltransferases secretion is sensitive to the γ-secretase inhibitor DFK-167 but is independent of Presenilins. Cells expressing ST6Gal-I were treated for 24h at 37°C with the following inhibitors, the β-secretase inhibitor C3, the γ-secretase inhibitor DFK-167 or the metalloprotease inhibitor GM6001. Inhibitor stocks were made in DMSO and added to cells in serum-free OptiMEM medium at 1% final concentration of DMSO. Culture media that were not treated with inhibitors were supplemented with 1% DMSO. Proteins in media (M) were compared to their cell-bound counterparts (C) in the absence (-) or the presence (+) of the inhibitor. **A**) Effect of protease inhibitors on ST6Gal-I secretion. **B**) The experiment with DFK-167 was repeated and the Western blot was intentionally overexposed to emphasize the effect of the inhibitor. **C**) Effect of DFK-167 on the other GTs. Note that following DFK-167 treatment, some GTs are almost absent from cell culture media (*i.e* β1,3GnT-III and C2GnT-I). **D**) MEFs (WT) or those where Presenilin-1 and Presenilin-2 genes where deleted (PS-dKO) expressing either C2GnT-I, ST6Gal-I, FucT-I or GnT-V were analyzed for protein secretion as indicated above. Note that no difference in protein expression and secretion could be seen between cells (C) and media (M) whether Presenilin genes are present (WT) or absent (PS-dKO).

### Secretion of glycosyltransferases is not impared in Presenilin1 and Presenilin2-deficient fibroblasts

Using Presenilin1-deficient MEF cells, Nakahara and co-workers also showed that the DFK167-sensitive γ-secretase that released GnT-V is Presenilin-1 [17]. To ascertain whether this could be extended to other GTs, we obtained Presenilin1/2-deficient MEF cells from the same laboratory as Nakahara’s group; that is De Strooper’s laboratory (Katholieke Universiteit, Leuven and Flanders Institute for Biotechnology, Belgium) and transduced them with lentiviral constructs of gfp-tagged C2GnT-I, ST6Gal-I and FucT-I or FLAG-tagged GnT-V (used as a control). As shown in Fig. 2D, the amounts of secreted proteins (M) from wild-type (WT) or from Presenilin1/2-deficient cells (PS-dKO) are similar, suggesting that neither Presenilin-1 nor Presenilin-2 are likely to be involved in the shedding of these molecules. Regarding GnT-V, the soluble FLAG-tagged protein from WT migrates with apparent molecular weight of 90 kDa while cell-associated counterparts exhibit sizes of 100 and 75 kDa (Fig. 2D) which is consistent, at least in part, with what has been found previously by Nakahara and coworkers [17]. However, the 90 kDa released species are recovered whether or not Presenilins are present, indicating that the GnT-V may not be processed by these γ-proteases. Taken as a whole the above data suggest that GTs might be processed by DFK167-sensitive proteases, probably belonging to the i-CLiP proteases but different from Presenilins

The discrepancies between our present data and those obtained by Kitazume and coworkers on ST6Gal-I [16,22,35] on the one hand, and on GnT-V by Nakahara and colleagues [17] on the other hand, are incomprehensible, particularly because we used the same cell system [BACE1 KO, BACE1/2 dKO and Presenilin (PS) dKO] originating from the same laboratory (*i.e* the De Strooper’s group) and the same GTs, ST6Gal-I and GnT-V, aside from the fact that their studies were based on intact proteins while we herein used gfp or FLAG-tagged ones. Besides, it is worth noting that both BACEs and Presenilins are type-I membrane-bound proteins and therefore, would preferentially act on type-I membrane proteins (reviewed in [40]). Now, glycosyltransferases are type-II membrane proteins and therefore, should well be substrates of proteases having the same topological orientation. This is the case of the signal peptide peptidase (SPP) or the signal peptide peptidase-like (SPPL) which belong to the i-CLiP family and to the same GxGD gamma-secretase group as Presenilins, while having their YD and GxGD-containing domains inverted relative to Presenilins. Thus, these proteases exhibit active sites oriented in a consistent topology relative to their type-II transmembrane protein substrates (reviewed in [41]), such as glycosyltransferases,

### Some, but not all glycosyltransferases are released by the signal peptide peptidase (SPP)-like protease SPPL3

We therefore postulated that GTs could be processed by SPP/SPPL protease family. In this regard, Voss and co-workers have recently demonstated that the shedding of glycosyltransferases is rather mediated by the signal peptide peptidase SPPL3; the loss of which directly influenced the glycosylation state of numerous glycoprotein substrates [36]. As for the studies described above, we sought to use cells devoid of SPPL3 gene (SPPL3 KO MEF cells), but could not obtain such cell lines from laboratories that developed them. Instead, we merely established MEFs with altered SPPL3 expression by a CRISPR/Cas9-mediated genome-editing approach. As shown in Fig. 3, Western blotting with the anti-SPPL3 antibody reveals the lack of SPPL3 in cells that co-express Cas9 and the SPPL3- sepcific RNA guide (lane +), whereas a 37 and and 62 KDa species are detected in in control cells (lane -), that may correspond to monomer and homodimer entities of the mouse SPPL3, which is consistent with data reported by others[42] [43]. These Cas9/gRNA-treated MEFs were then transduced with lentiviral constructs of gfp-tagged C2GnT-I, β1,3GnT-III, ST6Gal-I, or FLAG-tagged GnT-V (again used as a control) and secreted proteins were analyzed as described above. As shown in Fig. 3B, the amounts of secreted proteins from intact cells (lane -) and Cas9/gRNA-treated ones (lane +) were similar for C2GnT-I and ST6Gal-I but dramaticaly lowered for GnT-V and were even completely absent in cell culture media of β1,3GnTIII-expressing cells. This suggests that SPPL3 may selectively act on some GTs and partially or not at all, on others.

**Figure 3:**
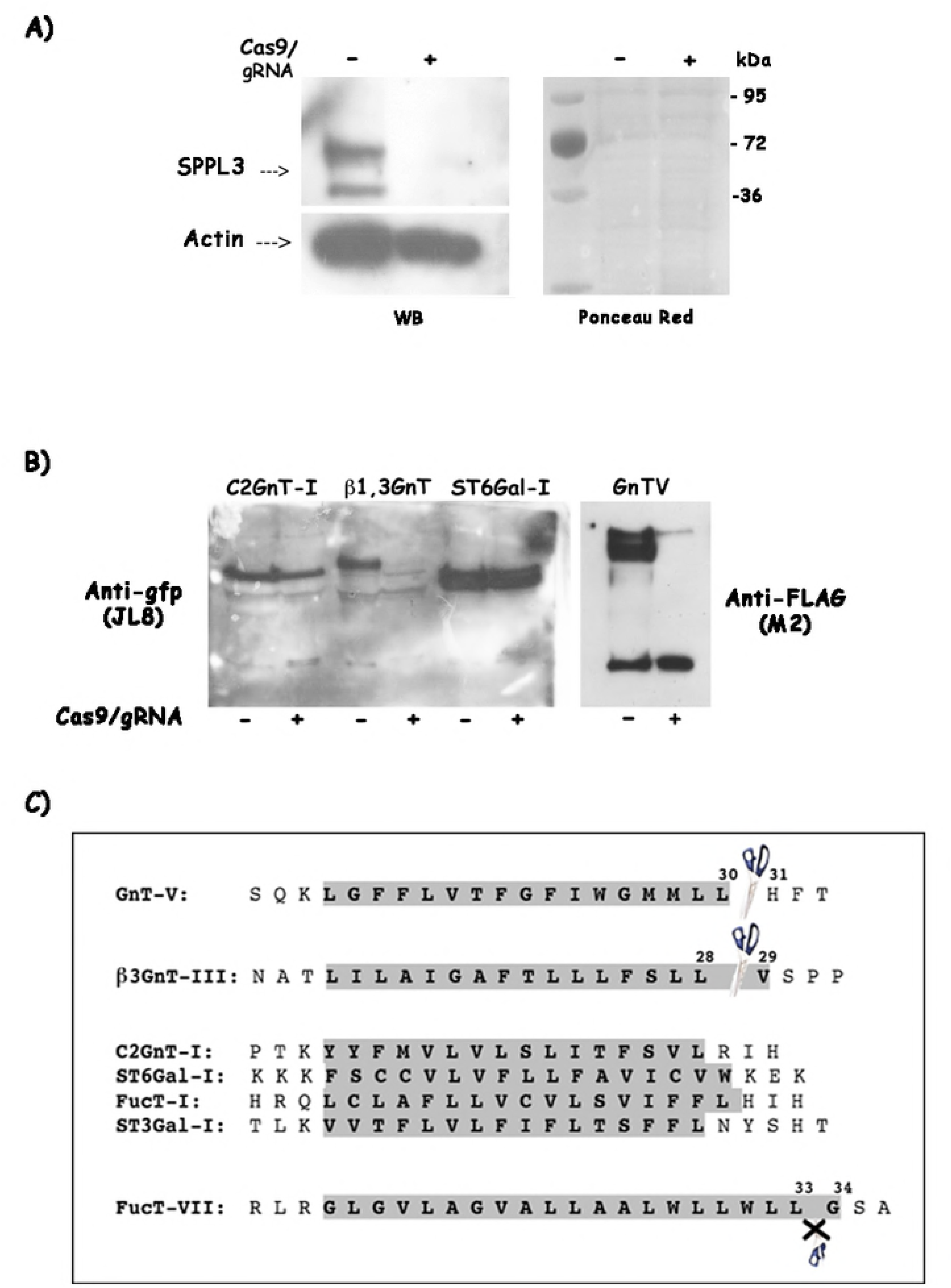
Suppression of SPPL3 activity in MEFs impairs shedding of some but not all GTs. **A)** Analysis of SPPL3 expression in control MEFs (-) and CRISPR/Cas9-treated cells (+). MEFs express SPPL3 as two proteins of 37 and 67 kDa whereas cells exposed to Cas9/gRNA completely lack the protease, while Ponceau Red staining confirms that equal amounts of protein were loaded. **B**) Western blots of culture media from GTs-expressing cells transduced (+) or not (-) by CRISPR/Cas9 lentiviral particles (Cas9/gRNA). Note that the secretion of β1,3GnT-III and GnT-V were dramatically impaired while C2GnT-I and ST6Gal-I were released normally. C) Schematic presentation of amino acid sequences of the GTs studied herein and their putative cleavage sites by SPPL3 (shown by scissors).

Once again, this result does not completely match the previously reported data on SPPL3 substrates [21,36] and rises the questions of why SPPL3 exerts such a selectivity towards glycosyltransferases. One would attempt a hypothesis that GnT-V and β3GnT-III are cleaved by SPPL3 because they share the same enzymatic property of transferring a GlcNAc residue on their cognate substrates. But then why C2GnT-I, which belongs to the same family of “branching GlcNAc-transferase” as GnT-V, is not processed by SPPL3? Secondly, why FucT-VII is not released from any cell model studied so far, while having the same dipeptidyl Leu-Leu at the lipid bilayer boundary, comparable to that of GnT-V [17] and β1,3GnT-III (see Fig. 3C). In fact, GnT-V has been shown to be cleaved between the double Leucine (Leu29Leu30) and His31 residues [17,44] and we herein demonstrate that β1,3GnT- III, the transmembrane domain of which also countains a double Leucine motif (Leu27Leu28) at the lipid bilayer boundary, is not shed from cells with impaired SPPL3 expression (Fig. 3C), suggesting that this GT might likewise be cut by SPPL3 at an equivalent position as GnT-V. These results also suggest that SPPL3 has a preference for a double leucine motif at the edge of the transmembrane domain and the cytosol. Hence, based on sequence specificity, one would expect FucT-VII to be potentially a SPPL3 substrate, since its transmebrane domain also ends with a double Leucine motif (see Fig. 3C). However, previous and present studies, have shown that FucT-VII has never been found in culture media [19].

### A right subcellular distribution is a prerequisite for glycosyltransferase proteolytic cleavage

With all these questions in mind, we reasoned that SPPL3 selectivity may not rely solely on a peptide sequence specificity, but other factors may contribute for the release of a given GT by SPPL3. We then decided to compare the intracellular distribution of SPPL3 *versus* C2GnT-I, β1,3GnT-III, ST6Gal-I and FucT-VII. The choice of FucT-VII was dictated by the followings i) as stated above, it is the only uncleaved protein among our GTs, ii) it generally exhibits a broad and diffuse distribution rather than the typical perinuclear shape indicative of Golgi localization [8,33]. Unfortunately, only few data exist on SPPL3 intracellular distribution, except one study that localized the protease to the ER [43] and another which suggested a Golgi localization [36]. To address this issue, we used a mCherry-tagged version of SPPL3 [29] that we mounted in the lentiviral vector pL1 to transduced MEFs expressing gfp-tagged GTs. As shown in Fig. 4, confocal examination discloses, as expected, a typical Golgi staining for gfp-tagged C2GnT-I and ST6Gal-I, which is clearly distinguishable from that of mCherry-SPPL3 (Fig. 4A and 4C). However, a colocalization between β1,3GnTIII-gfp and mCherry SPPL3 could be seen in some cells at the juxtanuclear areas that may correspond to Golgi cisternae [orange color (arrow), Fig. 4B). FucT-VII distribution however, is clearly distingushable from that of mCherry SPPL3 (Fig. 4 D). This result is to be paralleled with the above data and provide a possible explanation of i) why β1,3GnTIII is released by SPPL3 whereas C2GnT-I and ST6Gal-I are not and ii) why FucT-VII is not released while having a putative SPPL3-cleavable sequence in its transmembrane domain, but does not colocalize with the latter. Therefore, we postulate that the cleavage efficiency of GTs by SPPL3 requires not only a specific sequence but also an adequate intracellular distribution.

**Figure 4:**
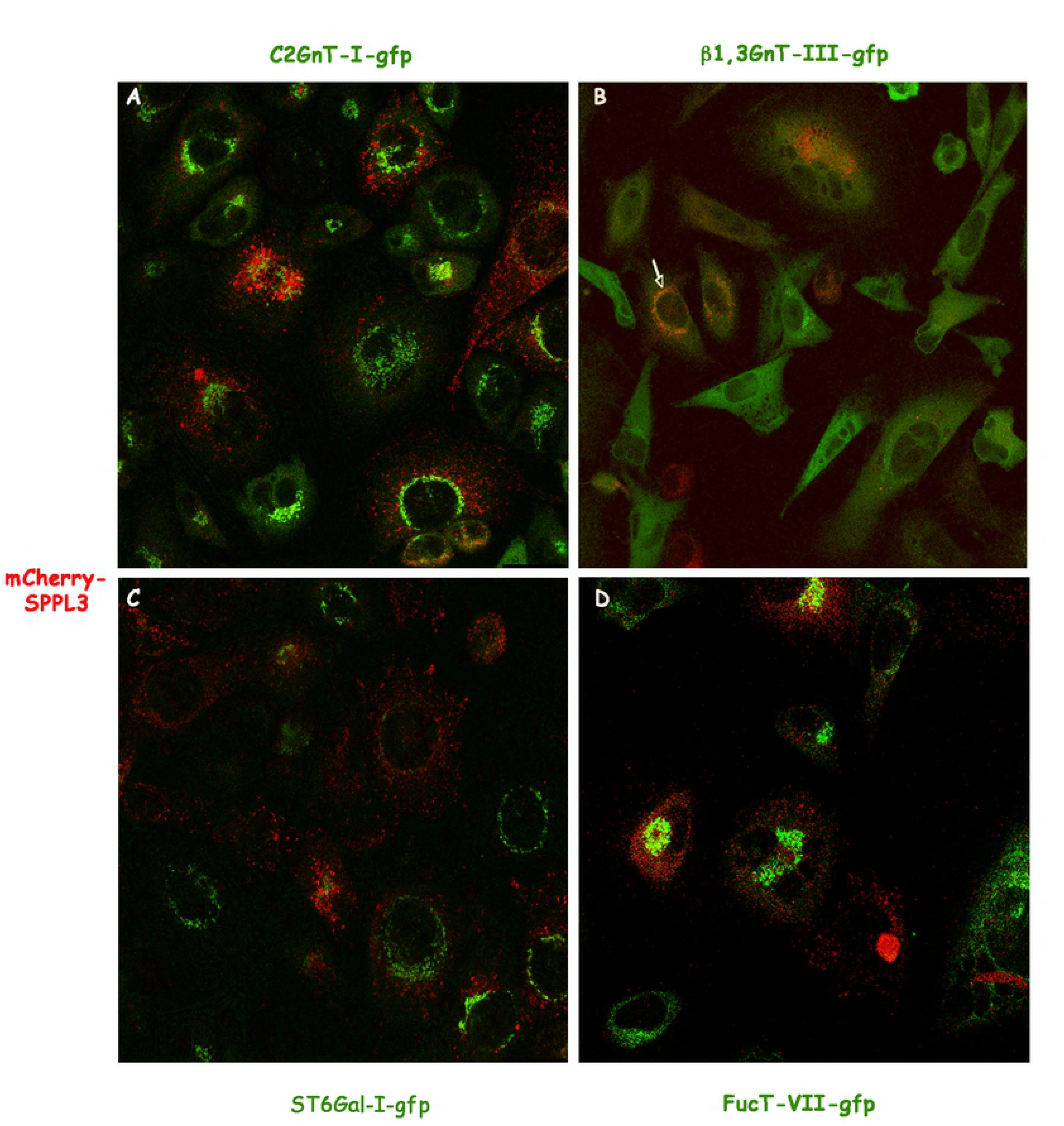
Intracellular distribution of GTs-gfp (green) and mCherry-SPPL3 (red). MEFs expressing gfp-tagged C2GnT-I, β1,3GnT-III, ST6Gal-I or FucT-VII were transduced with pL1 lentiviral vector harboring mCherry-SPPL3. Living cells were then examined by confocal microsopy as indicated under Material and Methods. Images were processed with Metamorph Imaging system version 3.5 and transferred to Adobe Photoshop as TIFF files.

### Displacing FucT-VII to the β1,3GnT-III compartrment enhances its proteolytic cleavage and shedding

To furher ascertain the above data, we sought to move FucT-VII to the β1,3GnT-III compartment and by doing so, we wanted to know whether FucT-VII would become cleavable by SPPL3. For this purpose, we used the same strategy as in Zerfaoui *et al* [8], that is by fusing a protein portion to the transmembrane domain of a given GT, we were able to target this portion to the same subcellular compartment as the GT [8]. We therefore fused the first 29 amino acid sequence of β1,3GnT-III (encompassing the cytosolic and the transmembrane domain), to the catalytic domain of FucT-VII. The resulting chimeric enzyme (referred to as β1,3GnT-III/FucT-VII) was then transiently expressed in MEFs as gfp-tagged protein and compared to FucTVII-gfp, with respect to localization and secretion. Results are shown on Fig. 5. Confocal examination discloses, as expected, a diffuse distribution for FucT-VII-gfp, whereas the chimeric β1,3GnT-III/FucT-VII-gfp predominantly overlaps (yellow staining) with the medial-Golgi marker α-mannosidase-II (α-Man-II, red staining), indicative of a medial-Golgi localization (Fig. 5A). Next, the chimeric β1,3GnT-III/FucT-VII was stably expressed in MEFs by lentiviral transduction and compared to FucTVII-expressing cells with respect to released proteins. Interestingly, the mislocalized chimeric FucT-VII becomes now susceptible to proteolysis and a soluble protein could be detected in the medium (Fig. 5B, lane M). It is noteworthy that the size of the released fragment is closely similar to its cellular counterpart (lane C), a situation that recalls other shed GTs suggesting that the chimeric β1,3GnT-III/FucT-VII might be acted-up by the same protease as the one that cleaves β1,3GnT-III, presumably SPPL3.

**Figure 5:**
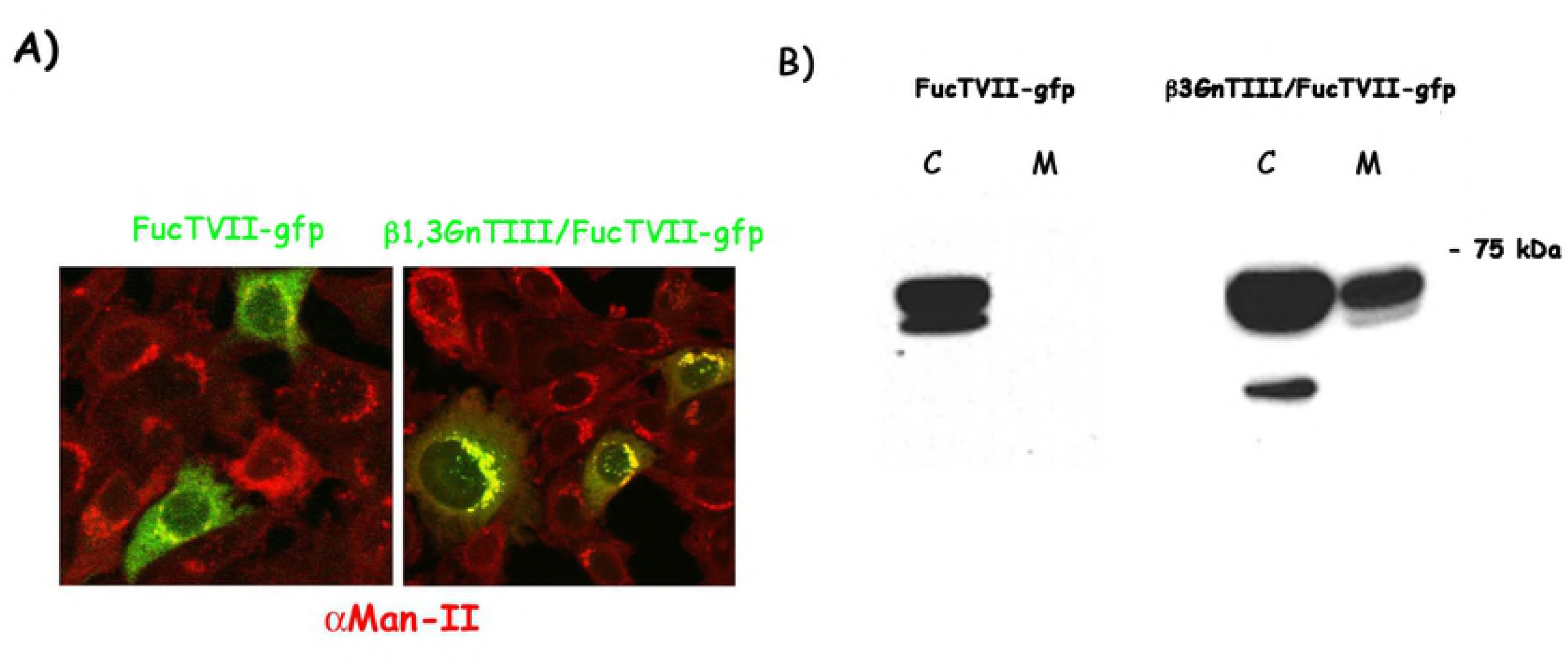
Analyses of intracellular distribution and secretion of intact FucTVII-gfp and the chimeric β1,3GnTIII/FucTVII-gfp. **A)** MEFs were transiently transfected with either FucTVII-gfp or the chimeric β1,3GnTIII/FucTVII-gfp constructs (green) and examined by confocal microscopy as above, with respect of the Golgi marker α-mannosidase-II (αMan-II, red). Note that the chimeric β1,3GnTIII/FucTVII-gfp overlaps very well with αMan-II (yellow, right panel), whereas FucTVII-gfp does not. **B)** Media from cells expressing FucTVII-gfp and those expressing the chimeric fucosyltransferase (β1,3GnTIII/FucTVII-gfp) were then collected and analyzed for FucT-VII secretion. Note that following FucT-VII displacement to β1,3GnTIII location caused its susceptibility to proteolytic cleavage and release in the medium.

In conclusion, we herein explored the possibility of Alzheimer disease-associated proteases, namely BACE-1 and BACE-2 or Presenilin-1 and Presenilin-2, and demonstrated that they do not play any role in the shedding of GTs. On the contrary, we confirm that SPPL3 could cleave some but not all glycosyltransferases, with a selectivity that we attribute to sequence specificity together with the probability of meeting between the protease and its cognate substrates along the Golgi stacks.

## Aknowledgements

The authors thank F. Silvy for assistance with confocal microscopy and Dr J. Luis for critically reading the manuscript. This work was supported by the Institut National de la Santé et de la Recherche Médicale (INSERM).

## Abbreviations

MEFs: mouse embryonic fibroblasts
BACE: beta-site amyloid precursor protein-cleaving enzyme
C2GnT-I: core 2 beta1,6-N-acetylglucosaminyltransferase-I
β3GnT-III: core 1 beta1,3-N-acetylglucosaminyltransferase-III
GnT-V: beta1,6-N-acetylglucosaminyltransferase-V
FucT-I: alpha1,2-fucosyltransferase-I
FucT-VII: alpha1,3- fucosyltransferase-VII
ST3Gal-I: alpha2,3-sialyltransferase-I
ST6Gal-I: alpha2,6- sialyltransferase-I

## REFERENCES

1. Sperandio M, Gleissner CA, Ley K (2009) Glycosylation in immune cell trafficking. Immunol Rev 230: 97–113.

2. Rabinovich GA, Croci DO (2012) Regulatory circuits mediated by lectin-glycan interactions in autoimmunity and cancer. Immunity 36: 322–335.

3. Cui H, Freeman C, Jacobson GA, Small DH (2013) Proteoglycans in the central nervous system: role in development, neural repair, and Alzheimer’s disease. IUBMB Life 65: 108–120.

4. Ouwerkerk JP, de Vos WM, Belzer C (2013) Glycobiome: bacteria and mucus at the epithelial interface. Best Pract Res Clin Gastroenterol 27: 25–38.

5. Banerjee DK (2012) N-glycans in cell survival and death: cross-talk between glycosyltransferases. Biochim Biophys Acta 1820: 1338–1346.

6. Lairson LL, Henrissat B, Davies GJ, Withers SG (2008) Glycosyltransferases: structures, functions, and mechanisms. Annu Rev Biochem 77: 521–555.

7. Opat AS, Houghton F, Gleeson PA (2001) Steady-state localization of a medial-Golgi glycosyltransferase involves transit through the trans-Golgi network. Biochem J 358: 33–40.

8. Zerfaoui M, Fukuda M, Langlet C, Mathieu S, Suzuki M, et al. (2002) The cytosolic and transmembrane domains of the beta 1,6 N-acetylglucosaminyltransferase (C2GnT) function as a cis to medial/Golgi-targeting determinant. Glycobiology 12: 15–24.

9. Jackson CL (2009) Mechanisms of transport through the Golgi complex. J Cell Sci 122: 443–452.

10. Tu L, Banfield DK (2010) Localization of Golgi-resident glycosyltransferases. Cell Mol Life Sci 67: 29–41.

11. Paulson JC, Colley KJ (1989) Glycosyltransferases. Structure, localization, and control of cell type-specific glycosylation. J Biol Chem 264: 17615–17618.

12. Weinstein J, Lee EU, McEntee K, Lai PH, Paulson JC (1987) Primary structure of beta-galactoside alpha 2,6-sialyltransferase. Conversion of membrane-bound enzyme to soluble forms by cleavage of the NH2-terminal signal anchor. J Biol Chem 262: 17735–17743.

13. Masri KA, Appert HE, Fukuda MN (1988) Identification of the full-length coding sequence for human galactosyltransferase (beta-N-acetylglucosaminide: beta 1,4-galactosyltransferase). Biochem Biophys Res Commun 157: 657–663.

14. Homa FL, Hollander T, Lehman DJ, Thomsen DR, Elhammer AP (1993) Isolation and expression of a cDNA clone encoding a bovine UDP-GalNAc:polypeptide N-acetylgalactosaminyltransferase. J Biol Chem 268: 12609–12616.

15. Jaskiewicz E, Zhu G, Bassi R, Darling DS, Young WW Jr., (1996) Beta1,4-N-acetylgalactosaminyltransferase (GM2 synthase) is released from Golgi membranes as a neuraminidase-sensitive, disulfide-bonded dimer by a cathepsin D-like protease. J Biol Chem 271: 26395–26403.

16. Kitazume S, Suzuki M, Saido TC, Hashimoto Y (2004) Involvement of proteases in glycosyltransferase secretion: Alzheimer’s beta-secretase-dependent cleavage and a following processing by an aminopeptidase. Glycoconj J 21: 25–29.

17. Nakahara S, Saito T, Kondo N, Moriwaki K, Noda K, et al. (2006) A secreted type of beta1,6 N-acetylglucosaminyltransferase V (GnT-V), a novel angiogenesis inducer, is regulated by gamma-secretase. FASEB J 20: 2451–2459.

18. Larsen RD, Ernst LK, Nair RP, Lowe JB (1990) Molecular cloning, sequence, and expression of a human GDP-L-fucose:beta-D-galactoside 2-alpha-L-fucosyltransferase cDNA that can form the H blood group antigen. Proc Natl Acad Sci U S A 87: 6674–6678.

19. El-Battari A, Prorok M, Angata K, Mathieu S, Zerfaoui M, et al. (2003) Different glycosyltransferases are differentially processed for secretion, dimerization, and autoglycosylation. Glycobiology 13: 941–953.

20. Saito T, Miyoshi E, Sasai K, Nakano N, Eguchi H, et al. (2002) A secreted type of beta 1,6- N-acetylglucosaminyltransferase V (GnT-V) induces tumor angiogenesis without mediation of glycosylation: a novel function of GnT-V distinct from the original glycosyltransferase activity. J Biol Chem 277: 17002–17008.

21. Kuhn PH, Voss M, Haug-Kroper M, Schroder B, Schepers U, et al. Secretome analysis identifies novel signal Peptide peptidase-like 3 (Sppl3) substrates and reveals a role of Sppl3 in multiple Golgi glycosylation pathways. (2015) Mol Cell Proteomics 14: 1584–1598.

22. Kitazume S, Tachida Y, Oka R, Shirotani K, Saido TC, et al. (2001) Alzheimer’s beta-secretase, beta-site amyloid precursor protein-cleaving enzyme, is responsible for cleavage secretion of a Golgi-resident sialyltransferase. Proc Natl Acad Sci U S A 98: 13554–13559.

23. Dominguez D, Tournoy J, Hartmann D, Huth T, Cryns K, et al. (2005) Phenotypic and biochemical analyses of BACE1- and BACE2-deficient mice. J Biol Chem 280: 30797–30806.

24. Herreman A, Hartmann D, Annaert W, Saftig P, Craessaerts K, et al. (1999) Presenilin 2 deficiency causes a mild pulmonary phenotype and no changes in amyloid precursor protein processing but enhances the embryonic lethal phenotype of presenilin 1 deficiency. Proc Natl Acad Sci U S A 96: 11872–11877.

25. Herreman A, Van Gassen G, Bentahir M, Nyabi O, Craessaerts K, et al. (2003) gamma-Secretase activity requires the presenilin-dependent trafficking of nicastrin through the Golgi apparatus but not its complex glycosylation. J Cell Sci 116: 1127–1136.

26. Mali P, Yang L, Esvelt KM, Aach J, Guell M, et al. (2013) RNA-guided human genome engineering via Cas9. Science 339: 823–826.

27. Saito H, Nishikawa A, Gu J, Ihara Y, Soejima H, et al. (1994) cDNA cloning and chromosomal mapping of human N-acetylglucosaminyltransferase V+. Biochem Biophys Res Commun 198: 318–327.

28. Makowski SL, Wang Z, Pomerantz JL A protease-independent function for SPPL3 in NFAT activation. (2014) Mol Cell Biol 35: 451–467.

29. Hamblet CE, Makowski SL, Tritapoe JM, Pomerantz JL NK Cell Maturation and Cytotoxicity Are Controlled by the Intramembrane Aspartyl Protease SPPL3. (2016) J Immunol 196: 2614–2626.

30. Wiznerowicz M, Trono D (2003) Conditional suppression of cellular genes: lentivirus vector-mediated drug-inducible RNA interference. J Virol 77: 8957–8961.

31. Mathieu S, Prorok M, Benoliel AM, Uch R, Langlet C, et al. (2004) Transgene expression of alpha(1,2)-fucosyltransferase-I (FUT1) in tumor cells selectively inhibits sialyl-Lewis x expression and binding to E-selectin without affecting synthesis of sialyl-Lewis a or binding to P-selectin. Am J Pathol 164: 371–383.

32. Laemmli UK (1970) Cleavage of structural proteins during the assembly of the head of bacteriophage T4. Nature 227: 680–685.

33. Prorok-Hamon M, Notel F, Mathieu S, Langlet C, Fukuda M, et al. (2005) N-glycans of core2 beta(1,6)-N-acetylglucosaminyltransferase-I (C2GnT-I) but not those of alpha(1,3)-fucosyltransferase-VII (FucT-VII) are required for the synthesis of functional P-selectin glycoprotein ligand-1 (PSGL-1): effects on P-, L- and E-selectin binding. Biochem J 391: 491–502.

34. Hemming ML, Elias JE, Gygi SP, Selkoe DJ (2009) Identification of beta-secretase (BACE1) substrates using quantitative proteomics. PLoS One 4: e8477.

35. Kitazume S, Nakagawa K, Oka R, Tachida Y, Ogawa K, et al. (2005) In vivo cleavage of alpha2,6-sialyltransferase by Alzheimer beta-secretase. J Biol Chem 280: 8589–8595.

36. Voss M, Kunzel U, Higel F, Kuhn PH, Colombo A, et al. Shedding of glycan-modifying enzymes by signal peptide peptidase-like 3 (SPPL3) regulates cellular N-glycosylation. (2014) EMBO J 33: 2890–2905.

37. Wolfe MS (2009) gamma-Secretase in biology and medicine. Semin Cell Dev Biol 20: 219–224.

38. Rademacher TW, Parekh RB, Dwek RA (1988) Glycobiology. Annu Rev Biochem 57: 785–838.

39. Bierhuizen MF, Fukuda M (1992) Expression cloning of a cDNA encoding UDP-GlcNAc:Gal beta 1-3-GalNAc-R (GlcNAc to GalNAc) beta 1-6 GlcNAc transferase by gene transfer into CHO cells expressing polyoma large tumor antigen. Proc Natl Acad Sci U S A 89: 9326–9330.

40. Steiner H (2004) Uncovering gamma-secretase. Curr Alzheimer Res 1: 175–181.

41. Golde TE, Wolfe MS, Greenbaum DC (2009) Signal peptide peptidases: a family of intramembrane-cleaving proteases that cleave type 2 transmembrane proteins. Semin Cell Dev Biol 20: 225–230.

42. Nyborg AC, Herl L, Berezovska O, Thomas AV, Ladd TB, et al. (2006) Signal peptide peptidase (SPP) dimer formation as assessed by fluorescence lifetime imaging microscopy (FLIM) in intact cells. Mol Neurodegener 1: 16.

43. Krawitz P, Haffner C, Fluhrer R, Steiner H, Schmid B, et al. (2005) Differential localization and identification of a critical aspartate suggest non-redundant proteolytic functions of the presenilin homologues SPPL2b and SPPL3. J Biol Chem 280: 39515–39523.

44. Voss M, Schroder B, Fluhrer R Mechanism, specificity, and physiology of signal peptide peptidase (SPP) and SPP-like proteases. (2013) Biochim Biophys Acta 1828: 2828–2839.

